# RNA-Seq analysis and transcriptome assembly of *Salicornia neei* reveals a powerful system for ammonium detoxification

**DOI:** 10.1101/2021.09.07.458783

**Authors:** Mónica Díaz-Silva, Jonathan Maldonado, Pamela Veloso, Nicol Delgado, Herman Silva, José A. Gallardo

## Abstract

**Background:** *Salicornia neei* is a halophyte plant that has been proposed for use in the phytoremediation of the saline wastewater generated by land-based aquaculture, which usually contains elevated concentrations of ammonium resulting from protein metabolism. To identify the molecular mechanisms related to ammonium response, we analyzed the transcriptome of *S. neei* in response to growth in saline water containing 3 mM ammonium and the Michaelis–Menten ammonium removal biokinetics.

**Results:** The RNA sequencing generated a total of 14,680,108 paired-end reads from the control and stressed conditions. De novo assembly using the CLC Genomic Workbench produced 86,020 transcripts and a reference transcriptome with an N50 of 683 bp.

A total of 45,327 genes were annotated, representing 51.2% of the contig predicted from de novo assembly. As regards DEGs, a total of 9,140 genes were differentially expressed in response to ammonium in saline water; of these, 7,396 could be annotated against functional databases. The upregulated genes were mainly involved in cell wall biosynthesis, transmembrane transport and antiporter activities, including biological KEGG pathways linked to the biosynthesis of secondary metabolites, plant hormone signal transduction, autophagy, and nitrogen metabolism. In addiction, a set of 72 genes was directly involved in ammonium metabolism, including glutamine synthetase 1 (GLN1), glutamate synthase 1 (GLT1), and ferredoxin-dependent glutamate synthase chloroplastic (Fd-GOGAT). Finally, we observed that the ammonium uptake rate increased with increasing ammonium concentrations, and tended toward saturation in the range of 3 to 4 mM.

**Conclusion:** Our results support the hypothesis that an ammonium detoxification system mediated by glutamine and glutamate synthase was activated in *S. neei* when exposed to ammonium and saline water. The present transcriptome profiling method could be useful when investigating the response of halophyte plants to saline wastewater from land-based aquaculture.

## 1. Introduction

*Salicornia neei* has been proposed for the treatment of saline wastewater produced by land-based aquaculture effluents in South America [1]. *S. neei* is a herbaceous succulent hydrohalophyte that is abundantly distributed throughout much of the western coastline of the South Pacific [2]. Similar to other related species from Europe and North America, *S. neei* has attracted great interest due to its potential use as a leafy green food, because it contains significant amounts of nutrients and functional metabolites [3, 4]. Its potential to germinate and be cultivated in different salinity gradients and nitrogen concentrations has recently been evaluated in land-based white shrimp (*Litopenaeus vannamei*) farming systems [5, 6] and constructed wetlands (CWs) [1].

Land-based monoculture production systems use a large amount of artificial food rich in nitrogenous compounds, which are not fully utilized and may contaminate the culture water [7] or the environment if they are discarded without treatment. Although the accumulation of these compounds is variable and depends on both the species and the water treatment system involved [8], studies have shown that 133 kg of nitrogen (N) is released into the environment for each ton of harvested product [9]. The main nitrogenated compounds that are generated as waste are ammonia (NH_3_^+^) and ammonium (NH_4_^+^), which generally emerge as the final products of protein metabolism [10]. These compounds are found in equilibrium depending on the pH, temperature [11], and salinity [12]. The accumulation of ammonia and ammonium and other derived products, namely, nitrate (NO_3_^−^) and nitrite (NO_2_^−^), deteriorates water quality [13-15] and can generate eutrophication processes [16-20] and even encourage the development of diseases in cultivated organisms [21]. Therefore, taking advantage of nitrogenated compounds through the implementation of integrated marine aquaculture systems with halophyte plants has been proposed as a more sustainable alternative for the future development of land-based marine aquaculture [8, 22].

Nitrate and ammonium, the main forms of inorganic N that plants absorb, act as nutrients and signals affecting the growth and metabolism of the plant [23]. In general, the preference of plants for nitrate or ammonium depends on both the genotype and environmental variables such as soil pH and the availability of other nutrients [24].

Ammonium is a nutrient with qualities of rapid absorption but excessive accumulation in tissues; in plants, high concentrations can lead to symptoms of toxicity if it is the only source of N [25]. However, the addition of ammonium and nitrate together can be favorable for the plant, with a synergistic action [26]. In ammonium-tolerant plants, such as of the *Salicornia* genus, the NH_4_ ^+^ assimilation activity is higher compared to glycophytes. Ammonium stimulates its assimilation via the upregulation of several key enzymes involved in ammonium metabolism [27, 28]. After direct uptake or conversion from NH_3_^+^, NH_4_^+^ is assimilated to glutamine and glutamate via the activity of glutamine synthetase (GS), glutamate synthase (GOGAT), and glutamate dehydrogenase (GDH). The products of these pathways are required for the biosynthesis of other nitrogenous compounds [29]. These enzymes are part of two distinct pathways, the GS/GOGAT and GDH pathways. GS assimilates ammonium by catalyzing the amination of glutamate to form glutamine, while GOGAT catalyzes the reductive transfer of an amide-amino group from glutamine to 2-oxoglutarate, with the production of glutamate. GDH catalyzes the reversible amination of 2-oxoglutarate and ammonium to form glutamate [30].

In most glycophyte plants, highly saline soils disrupt metabolism, mainly due to decreased N uptake, altered activities of nitrate (NO_3_), and ammonium- (NH_4_^+^) assimilating enzymes, inducing changes in amino acid synthesis and increasing the activity of hydrolyzing enzymes such as RNase, DNase, and protease, which lead to macromolecule degradation [31]. However, halophytic plants have the enzymatic potential to synthesize GS, GOGAT, and GDH in order to assimilate ammonium under saline conditions [32]. These enzymes have been demonstrated to be powerful regulators of gene expression, and may be involved in diverse stress responsiveness [33]. For example, many halophytic plants accumulate large amounts of soluble nitrogenous compounds that are involved in salt tolerance, such as proline and the proline analog 4-hydroxy-N-methyl proline, glycine betaine, pinitol, myoinositol, mannitol, sorbitol, O-methylmucoinositol, and polyamines (PAs) [32, 34, 35].

To identify the molecular mechanisms of ammonium response, we conducted a transcriptomic analysis of *S. neei* in response to ammonium in saline water. It was expected that *S. neei* would have a gene response profile typical of halophyte plants when exposed to abiotic stressors of both ammonium and salinity.

## 2. Materials and methods

### 2.1. Plant material collection

*S. neei* wild plants were collected from the “Pullally” salt marsh (Papudo, Chile; latitude, 32°24’
s50′′S; longitude, 71°23’35.01”W) (Fig. 1) and transported to the facilities of Marine Farms Inc. (Laguna Verde, Chile, latitude 33° 6’ 36”S; longitude 71° 40’ 48” O) for propagation using cuttings. Stem cuttings were placed in an aquaponic cultivation system until they developed roots. Then, 180 rooted cuttings of *S. neei* were obtained from Marine Farms Inc. and transported to controlled growth chambers at the Pontificia Universidad Católica de Valparaíso (Valparaíso, Chile, latitude 33° 1’ 21” S, longitude 71° 37’ 57” W). The cuttings were acclimated for 7 days in hydroponic culture pots using UV-filtered seawater (600 mM NaCl) to remove suspended particles and microorganisms. The cuttings were reared in a 12/12 h (day/night) photoperiod cycle at a temperature of 20 °C and constant oxygenation, until being used in ammonium removal biokinetics or RNA-seq experiments. No extra nitrogen source was added to the filtered seawater during the acclimatization period, which had a basal concentration of 28 ± 6.5 µM L^−1^ of ammonium.

**Fig. 1.**
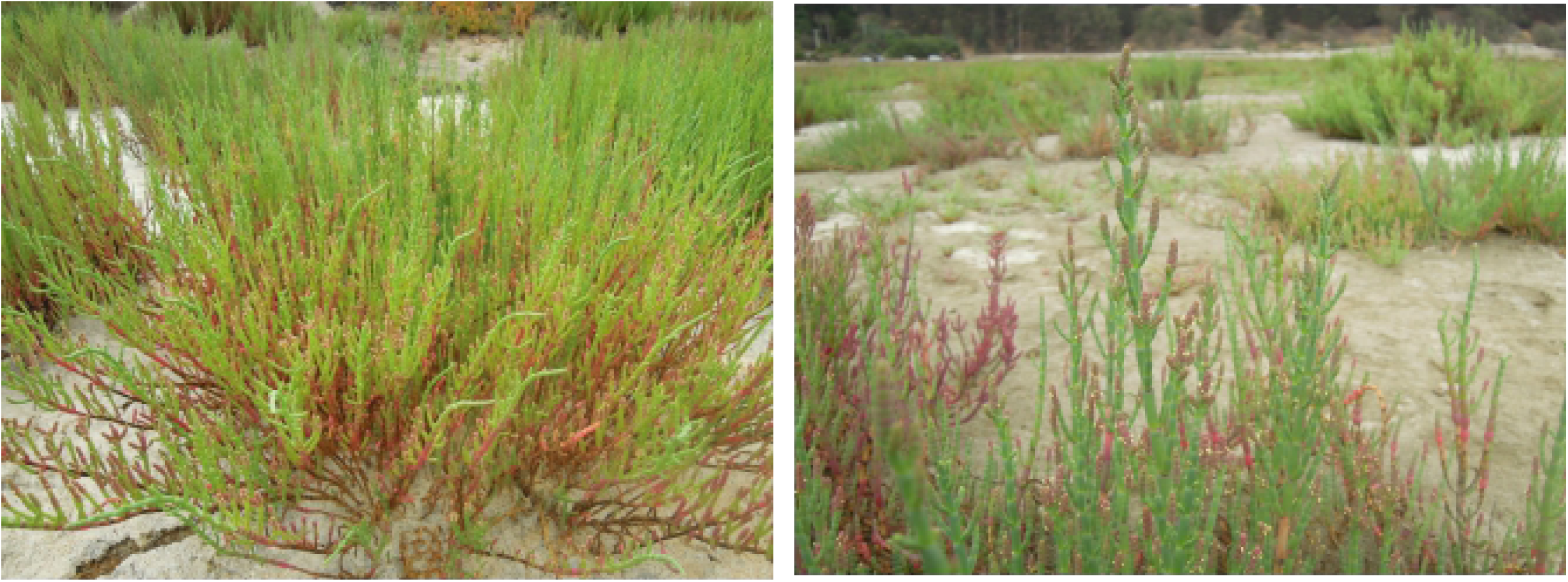
*Salicornia neei* wild plants from “Pullally” salt marsh (Papudo, Chile; latitude, 32°24’
s50′′S; longitude, 71°23’35.01”W).

### 2.2. Michaelis–Menten ammonium removal biokinetics

A total of sixty cuttings were used to characterize the Michaelis–Menten ammonium removal biokinetics. They were randomly distributed in five treatments that consisted of filtered seawater solutions with the addition of ammonium chloride (NH_4_Cl) at the following concentrations: 0, 1, 2, 3, and 4 mM. Each treatment was set up in 500 mL flasks in which 4 cuttings with an average weight of 21.8 ± 9.9 g were placed (Supplementary Fig. S1). For each treatment, a flask without plants was also used as a control or blank. Ammonium removal was evaluated every 60 min for a period of 5 h. At each time point, two 1.5 mL samples were taken from the treatments and control, and stored in Eppendorf tubes for subsequent analysis. The Nessler method [36] was used for the determination of ammonium, using a HACH DR2800 spectrophotometer at 425 nm. The ammonium uptake rate was expressed in µM NH_4_Cl g^−1^ DW h^−1^. After uptake, the root fresh weights were also recorded. The regressive relationship between uptake rate and ammonium concentration in the external solution was illustrated with the Michaelis–Menten [37] equation, as follows:

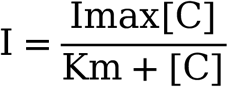

where I is uptake rate, Imax is the maximum uptake rate, Km is the half-saturation constant (Michaelis–Menten constant), and C is the ion concentration in solution. Michaelis–Menten curves were fitted using RStudio 7.0.

### 2.3. RNA-seq experiment and RNA extraction

A total of 24 cuttings were transferred to six plastic culture pots (four cuttings per pot and three biological replicates) containing 500 mL of filtered seawater with 0 or 3 mM NH_4_Cl. After 2.5 h, the cuttings were harvested for analysis. Approximately 1 g of collected stem tissue was extracted from each cutting, immediately frozen in liquid nitrogen, and stored at −80 °C until RNA extraction (Supplementary Fig. S2). Total RNA extraction was performed on three biological samples at 0 and 3 mM ammonium. RNA extractions were performed following the previously described pine tree extraction method [38] used in combination with β-mercaptoethanol [39] and TRIzol (Life Technologies, Corp., Carlsbad, USA) according to the manufacturer’s protocol. To homogenize the sample, it was ground in a mortar and pestle in the presence of liquid nitrogen and subsequently heated at 65 °C for 5 min in the presence of 1 mL pine tree buffer (PTB). Then, 20 µL β-mercaptoethanol was added to achieve greater sample purification, and the TRIzol steps were carried out according to the manufacturer’s protocol. Each sample was eluted in 60 μL of DEPC water and subjected to quantification by fluorometry using a Nanodrop (Biotek). Then, 5 µg of each sample was treated with RNase-free DNase I (Thermo Fischer Science) to remove residual genomic DNA, and precipitated with 3 M NaAc pH 5.2 and 100% EtOH. Finally, integrity was evaluated by electrophoresis in a 1% agarose gel, as described in [40].

### 2.4. Library construction, deep sequencing, and de novo transcriptome assembly

Library construction and deep sequencing of *S. neei* following treatment with 0 or 3 mM NH_4_Cl were performed at Macrogen (Inc. Seoul, South Korea) using the Solexa HiSeq2000 platform with the Truseq mRNA library previously constructed for paired-end applications, according to Macrogen’s protocol. For de novo transcriptome assembly, the raw data were cleaned of adaptor sequences, while low-quality reads (Q-value ≤ 20), reads with poly-N segments (reads containing more than 50% unknown bases) and short reads (less than 50 base pairs (bp)) were removed. The sequence quality algorithm contained in the CLC Genomics Workbench version 8.0 (http://www.clcbio.com) was used for this purpose. These results were processed using the scaffolding contig algorithm in the de novo assembly function of the CLC Genomics Workbench with default parameters. Single pools of 3 samples of 0 mM NH_4_Cl and 3 samples of mM NH_4_Cl were mapped separately against the assembled set of contigs using the RNA-seq tool of the CLC Genomics Workbench with the following parameters: similarity = 0.9; length fraction = 0.6; maximum mismatches = 2; unspecific match limit = 10. Paired reads were counted as 2, and paired-end distances were set as 101 bp.

### 2.5. Functional annotation

To identify the putative biological function, all assembled contigs were searched against public protein databases using the BLASTx algorithm of BLAST version 2.6.0+ [41]; the searched databases included the NCBI non-redundant (NR) protein database with an E-value cut-off of 10^−5^, the Kyoto Encyclopedia of Genes and Genomes (KEGG) pathway database with an E-value of 10^−5^, and EggNOG (orthology predictions and functional annotation) with an E-value of 10^−10^. Furthermore, to consolidate the information, all contigs were assigned to Gene Ontology (GO) categories using the Blast2Go version 5.2 [42] software package and an E-value cut-off of 10^−5^. Finally, because the genus *Salicornia* is poorly represented in the protein databases, contigs were also aligned against the nucleotide sequences of *Salicornia* available from the NCBI database using BLASTn with an E value of 10^−5^.

### 2.6. Differentially expressed genes (DEGs)

To determine the differentially expressed genes in *S. neei* under the two nutritional conditions, RNA-seq analysis was performed using the CLC Genomics Workbench program version 8.0 (http://www.clcbio.com). The relative transcript levels were defined as the number of reads that uniquely mapped to a gene. The expression levels were compared using a Z-Test [43-46] with 0 mM NH_4_Cl as the reference. This test compares counts by considering the proportions that make up the total sum of counts in each sample, correcting the data for sample size. For visual inspection, the original expression values were log10-transformed and then normalized using the quantile method that best fits the results [44]. As a significant threshold for the selection of genes with differential expression, a false discovery rate (FDR) p-value < 0.001 and an absolute fold change (FC) of 2.0 were adopted. Finally, GO enrichment analyses were performed separately on genes up-regulated in the 0 and 3 mM NH_4_Cl treatments to investigate the differences in the ammonium response mechanism.

## 3. Results

### 3.1. Kinetics of ammonium removal

The parameters of the NH_4_-N uptake kinetics have not been previously determined for *S. neei*. The ammonium uptake rate increased with increasing ammonium concentration in the external solution, and tended toward saturation in the range of 3 to 4 mM (Fig. 2). Using regression analysis, it was found that the kinetic characteristics of ammonium uptake by the test plant could be illustrated using the Michaelis–Menten equation at a significance level of p < 0.001. The kinetic parameters for ammonium uptake were estimated as a maximum rate (Imax) of 7.07 ± 0.27 and a half-saturation constant (Km) of 0.85 ± 0.12. Ammonium removal was significantly different from zero in all treatments (Supplementary Table S1).

**Fig. 2.**
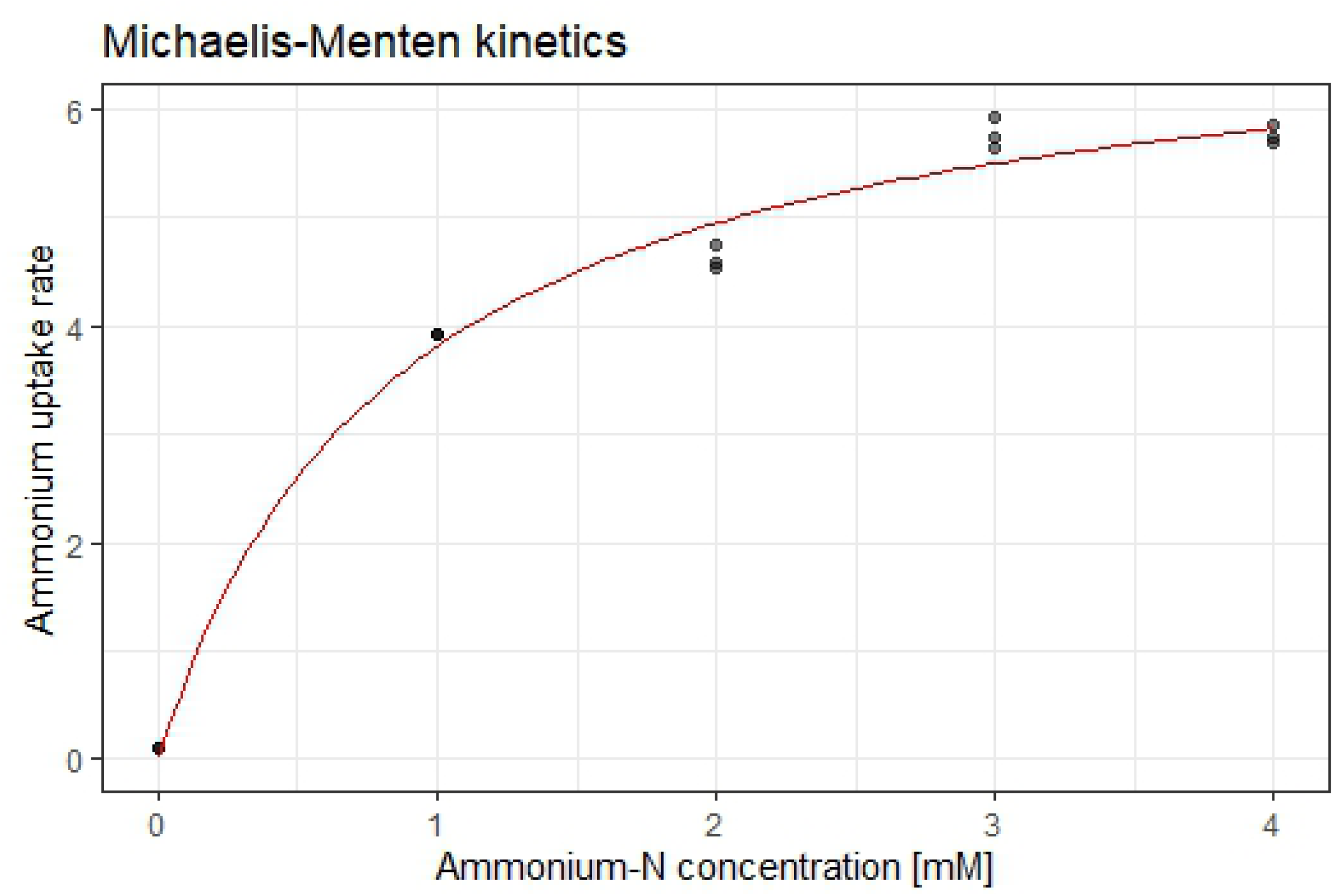
*S. neei* ammonium uptake rate as a function of ammonium concentration.

### 3.2. Transcriptomic sequencing, de novo assembly, and data availability

The transcriptomic analysis of pooled samples resulted in a total of 147,935,752 clean reads; the comprehensive reads were assembled into contigs using paired-end reads, resulting in 86,020 contigs with an average length of 586 bp (Table 1a and Table 1b). The raw sequencing data were deposited in the NCBI Short Read Archive (SRA) database under accession code SRR9694999, and they have been assigned the BioProject accession number PRJNA554118.

**Table 1a.** Reads trimming and mapping stats

|  |  | *Library-0mM-*<br>*NH<sub>4</sub>Cl* | *Library-3mM- NH<sub>4</sub>Cl* |
| --- | --- | --- | --- |
| **Quality and adapter trimming** | Number of reads | 71,793,586 | 74,986,522 |
|  | Avg length (bp) | 97,7 | 97,6 |
|  | Number of reads after trim | 71,695,733 | 74,879,764 |
| **Reads mapping to de novo assembled *S. neei* transcriptome** | Reads mapped in pairs | 51,430,428 | 51,493,796 |
|  | Reads mapped in broken pairs | 7,007,230 | 9,483,298 |
|  | Reads not mapped | 13,161,064 | 13,796,746 |
|  | Total | 71,598,722 | 74,773,840 |

**Table 1b.**
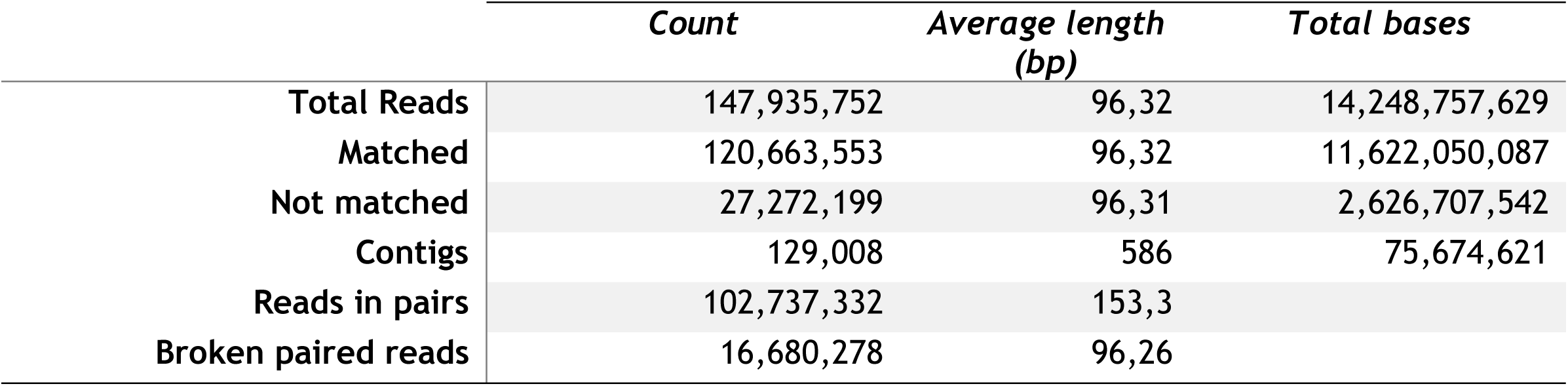
Summary of RNA-seq and de novo sequence assembly for *Salicornia neei*.

|  | *Count* | *Average length*<br>*(bp)* | *Total bases* |
| --- | --- | --- | --- |
| **Total Reads** | 147,935,752 | 96,32 | 14,248,757,629 |
| **Matched** | 120,663,553 | 96,32 | 11,622,050,087 |
| **Not matched** | 27,272,199 | 96,31 | 2,626,707,542 |
| **Contigs** | 129,008 | 586 | 75,674,621 |
| **Reads in pairs** | 102,737,332 | 153,3 |  |
| **Broken paired reads** | 16,680,278 | 96,26 |  |

### 3.3. Overview of sequences annotation and differentially expressed genes (DEGs)

The functional annotation process of the contigs using the NR database determined the presence of 86,020 contigs, most of which were assigned to *Spinacia oleracea* and *Chenopodium quinoa*. Most sequence homologies (Table 2) were found via comparisons against the NCBI–NR database (45,327 contigs), followed by GO (37,784 contigs) and EggNOG (20,047 contigs). Furthermore, 32,609 contigs were compared locally against the nucleotide sequences of *Salicornia* sp. Thus, a total of 13,906 new sequences from *S. neei* were found that did not match with those of other databases (supplementary Fig. S3). Based on the *S. neei* global transcriptome (86,020 contigs), 9,140 contigs were differentially expressed in their response to ammonium in saline water (7,040 up-regulated and 2,100 down-regulated), but only 7,396 could be annotated against the NR, GO, EggNOG and KEGG databases. Moreover, 588 DEGs were unique to the ammonium treatment. This was judged based on the abundance of transcription in the control, which was too low.

**Table 2.** Summary of annotations of assembled *Salicornia neei* contigs.

|  | *Number of contigs annotated* | *Percentage of contigs annotated* |
| --- | --- | --- |
| NCBI (NR) | 45,327 | 52.6 |
| NCBI (NT *Salicornia*) | 32,609 | 37.9 |
| GO | 37,784 | 43.9 |
| KEGG | 13,604 | 15.8 |
| EggNOG | 20,047 | 23.3 |

**GO:** Gene Ontology; **KEGG:** Encyclopedia of Genes and Genomes

### 3.4. Function annotation and classification

A total of 150,944 GO terms were assigned to 37,784 (43.9%) of the analyzed sequences; in multiple cases, several terms were assigned to the same sequence (mean = 4). Of the GO functional groups, 11 were assigned to biological processes, 10 to molecular functions, and 7 to cellular components. The top five functional groups were organic substance metabolic process (18,160), intracellular anatomical structure (18,042), cellular metabolic process (17,526), primary metabolic process (17,076), and organelle (15,242) (Fig. 3). GO terms retrieved for the DEGs of *S. neei* were subjected to a functional enrichment analysis, obtaining 17 enriched terms belonging to three main categories (BP, MF, CC) (Fig. 4). The enrichment proportion in the *S. neei* test set corresponds to the double or triple proportion of those found in the reference set: Xyloglucan metabolic process (0.3% in test set vs. 0.05% in reference set), beta-glucan metabolic process (0.72% in test set vs. 0.25% in reference set), beta-glucan biosynthetic process (0.63% in test set vs. 0.22% in reference set), cell wall organization biogenesis (1.25% in test set vs. 0.36% in reference set), sterol metabolic process (0.35% in test set vs. 0.10% in reference set), steroid metabolic process (0.46% in test set vs. 0.14% in reference set), glucosyltransferase activity (1.0% in test set vs. 0.37% in reference set), ion channel activity (0.7% in test set vs. 0.3% in reference set), and extracellular region (1.3% in test set vs. 0.69% in reference set) (see Fig. 4).

**Fig. 3.**
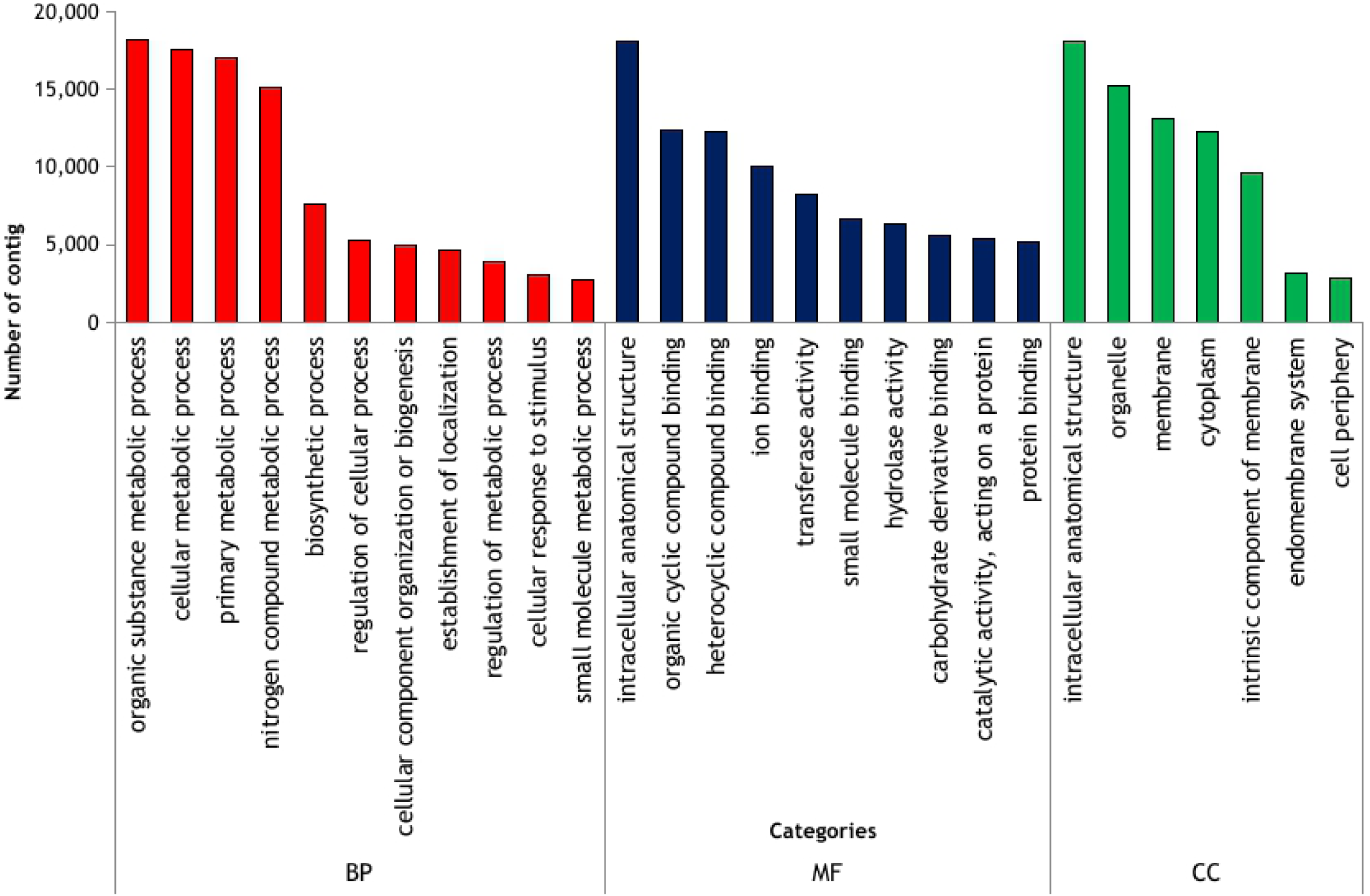
Histogram of Gene Ontology (GO) classification of 37,784 *S. neei* transcripts assigned to three main categories: biological process (BP), molecular function MF) and cellular component (CC).

**Fig. 4.**
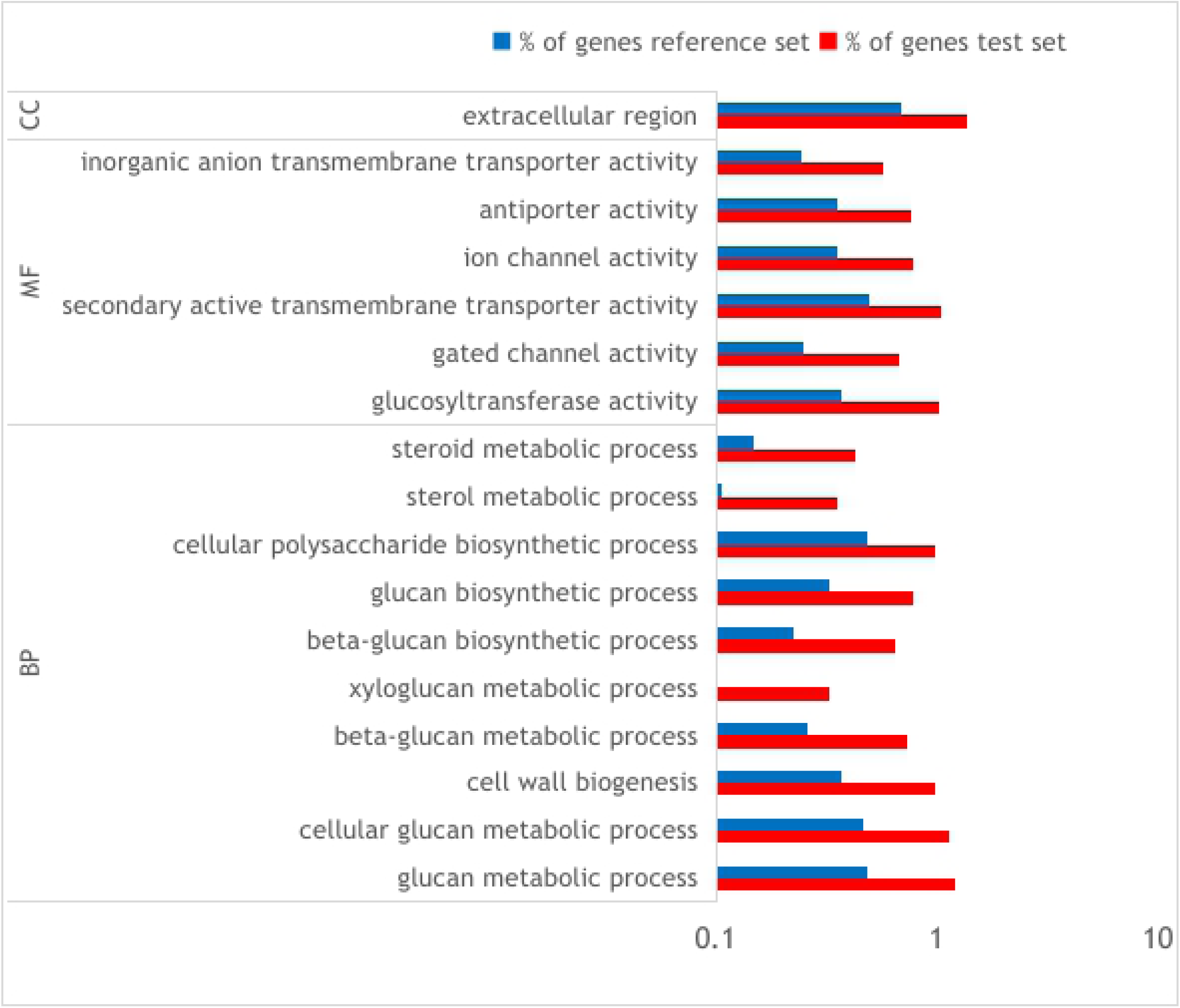
Gene Ontology enrichment analysis of differentially expressed genes in *S. neei* transcripts upregulated in test set vs upregulated genes in reference set.

Several DEGs were induced in response to ammonium treatment, mostly related to the maintenance of homeostasis. Out of the 91 DEGs related to nitrogen metabolism, 72 were up-regulated, including the following: glutamine synthetase (GS), 22-fold; oxoglutarate dehydrogenase (2-OGD), 2.34-fold; ferredoxin-dependent glutamate synthase chloroplastic (Fd-GOGAT), 4.29-fold; glutamate synthase 1 (NADH) chloroplastic isoform X1 (GLT1), 6.74-fold and ammonium transporter (AMT1), 2.39-fold (Supplementary Table S2). Phytohormone biosynthesis was also up-regulated; here, we found genes related to ethylene biosynthesis (S-adenosylmethionine synthetase (S-AdoMet) (20.82-fold), S-adenosyl-L-methionine (SAM) (4.8-fold), 1-aminocyclopropane-1-carboxylate synthase (ACC) (13.96-fold)) and abscisic acid biosynthesis (zeaxanthin epoxidase (ZEP) (2-fold), violaxanthin de-epoxidase, chloroplastic (VDE1) (2.56-fold) and xanthoxin dehydrogenase (ABA2) (2-fold)). Were also found many genes closely related to polyamine biosynthesis and autohagy, such as arginine decarboxylase-1 (ADC1), 2.36-fold; s-adenosylmethionine decarboxylase proenzyme (SAMDC1), 2.25-fold; polyamine oxidase-1 (PAO-1), 17.56-fold; autophagy-related protein 1 (ATG1), 2-fold; autophagy-related protein 13 (ATG13), 2.11-fold; autophagy-related protein 9 (ATG9), 5.89-fold; autophagy-related protein 2 (ATG2), 3.91-fold; and autophagy-related protein 18, 2.57-fold. Genes responsible for vesicle nucleation, such as vacuolar protein sorting 34 (VPS34) (3.26-fold) and vacuolar protein sorting 15 (VPS15) (2.0-fol), were also found up-regulated. Finally, genes responsible for the elongation and closure vacuolar protein sorting 8 (ATG8) were up-regulated 2.0-fold, and phosphatidylserine decarboxylase proenzyme 2 (PSD2) was up-regulated 2.0-fold; both of these catalyze the formation of phosphatidylethanolamine (PE).

DEGs were almost mapped to the metabolic pathway in KEGG; 1,206 DEGs were mapped to 167 KEGGs. Among the 167 pathways, the largest pathway was the biosynthesis of secondary metabolites (296 up-regulated genes), followed by ubiquitin-mediated proteolysis (55 up-regulated genes) and plant hormone signal transduction (48 up-regulated genes). DEGs were mostly mapped to autophagy (other (20 up-regulated genes) and nitrogen metabolism (14 up-regulated genes)) (see Table 3). Finally, to identify the proteins distributed in eukaryotic orthologous groups, and in clusters of orthologous groups and non-supervised orthologous groups, the annotated contigs were mapped to the annotations of the corresponding orthologous groups in the EggNOG database. A total of 20,047 (23.3%) sequences were divided into 25 categories and specific functional groups were identified (see Fig. 5). In the 25 EggNOG categories, the largest proportion of contigs belong to the cluster of “function unknown” (9,012; 44.9%), followed by “O: posttranslational modification, protein turnover, chaperones” (2,948; 14.7%) and “T: signal transduction mechanisms” (3,031; 15.11%). Other important groups closely related to ammonium nutrition response and stress response were “E: amino acid transport and metabolism” (1,391; 6.9%), “Q: Secondary metabolites biosynthesis, transport, and catabolism” (883; 4.4%), “M: cell wall/membrane/envelope biogenesis” (486; 2.42), and “V: defense mechanisms” (259; 1.2%). Only a small number of contigs were assigned to “W: extracellular structures” (6; 0.02%) or “Y: nuclear structure” (14; 0.006%).

**Table 3.** Summary of distribution of Kyoto Encyclopedia of Genes and Genomes (KEGG) pathways in the *S. neei* transcriptome.

|  | **KEGG**<br>**Pathway** | **N° Contigs** |
| --- | --- | --- |
| **Biosynthesis of secondary metabolite** | map01110 | 296 |
| **Ubiquitin mediated proteolysis** | map04120 | 55 |
| **Plant hormone signal transduction** | map04075 | 48 |
| **Endocytosis** | map04144 | 40 |
| **Starch and sucrose metabolism** | map00500 | 32 |
| **Circadian rhythm - plant** | map04712 | 24 |
| **Autophagy - other** | map04136 | 20 |
| **Arginine and proline metabolism** | map00330 | 18 |
| **Nitrogen metabolism** | map00910 | 14 |
| **Purine metabolism** | map00230 | 13 |

**Fig. 5.**
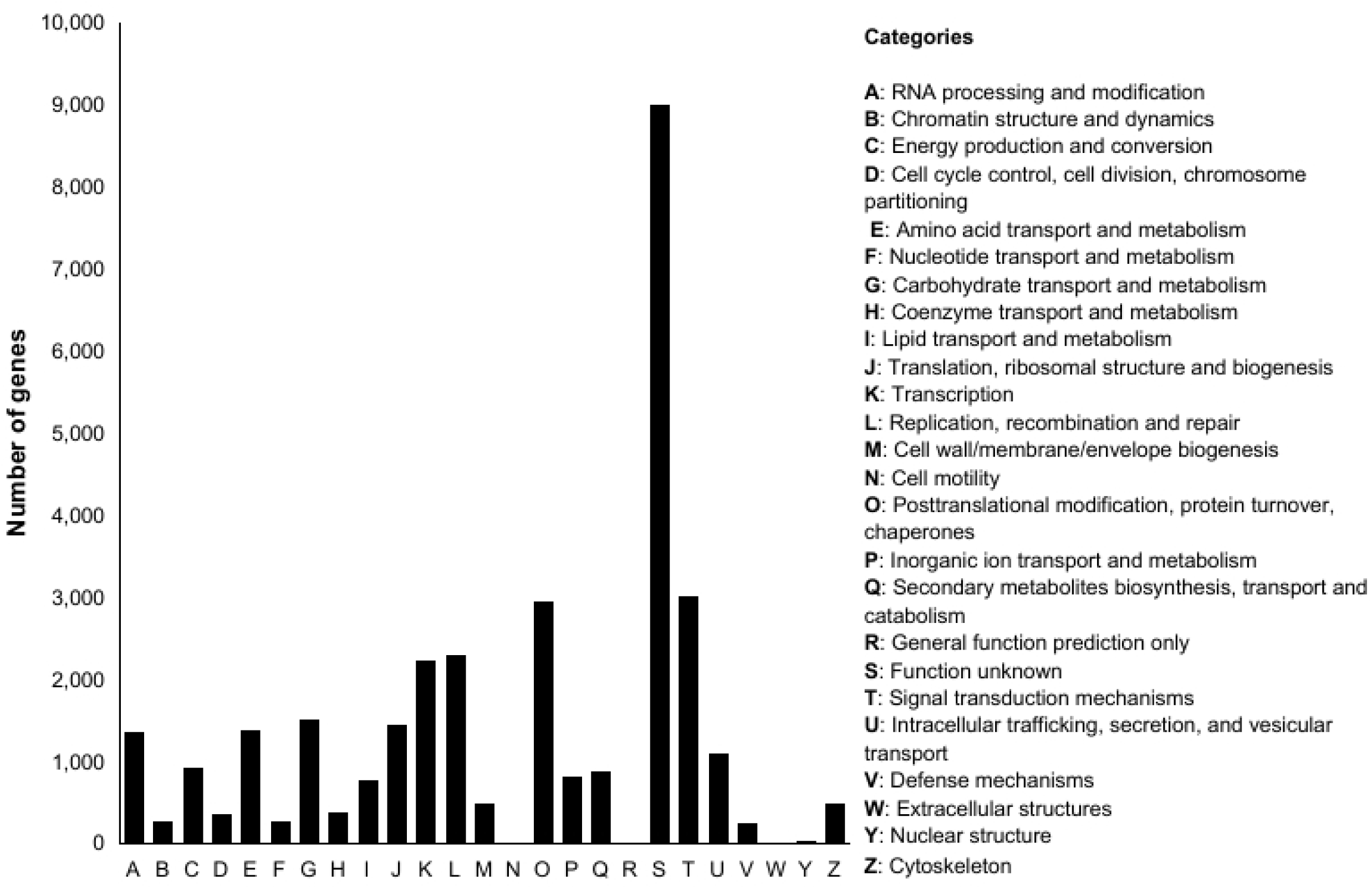
Functional annotation of contigs identified in *Salicornia neei* transcriptome. EggNOG classification analysis

## 4. Discussion

The use of halophyte plants to treat wastewater highly loaded with nitrogen compounds derived from the metabolism of aquatic organisms has been proposed to increase the sustainability of land-based aquaculture. However, the molecular mechanisms by which these plants can absorb and use these compounds in highly saline environments are poorly understood [20, 47]. This work used transcriptome analysis of the halophyte plant *S. neei* to investigate the possible pathways that are implicated in their response to ammonium nutrition. We considered possible ammonium homeostasis through ammonium metabolism and the encapsulation of ammonium in vacuoles to avoid increasing the acidity in the apoplast. We studied the kinetics of ammonium absorption in concentration gradients to demonstrate the activity of HATS (high-affinity transport systems) (AMT1), which are mainly related to the entry of ammonium into the cell, and therefore the disappearance of ammonium in the substrate.

The NH_4_ uptake kinetics of *S. neei* plants incubated at different concentrations fitted the Michaelis–Menten model well, up to 4 mM NH_4_ L^−1^ (Fig. 2), indicating the plant’s HATS activity at the tested concentrations. Quinta et al. [7] found a good fit for the Michaelis–Menten model up to 2 mM NH_4_ L^−1^ in *S. europeae*, and suggested that HATS (high-affinity transport systems) are responsible for N uptake in that plant. In this study, *S. neei* displayed higher ammonium absorption compared to *S. europeae* [7], demonstrating a better capacity for use in aquaculture wastewater.

The Michaelis–Menten equations obtained with the tested ammonium concentrations (1 to 4 mM NH_4_Cl) describe changes in ammonium absorption according to the amount of ammonia supplied. It was found that *S. neei* had a high affinity for the substrate (Km = 0.85 ± 0.12 mM NH_4_ L^−1^), indicating that this plant can perform well in substrates with high concentrations of ammonia. In addition, for aquaculture wastewater with concentrations equal to or greater than 4 mM NH_4_ L^−1^ and up to 7.07 ± 0.27 mM NH_4_ g^−1^ FW h^−1^, it is believed that the uptake rate does not increase because the plant blocks AMTs at higher concentrations. Concerning the kinetic characteristics of N uptake in plants, it has been proposed that species with higher maximum uptake rates might be better suited to cleaning up wastewater with high nutrient concentrations [48].

The cell wall plays an important role in stress perception by facilitating the activation of signaling pathways and remodeling growth strategies in response to stresses [49]. This structure constitutes the first line of defense against biotic and abiotic environmental influences through wall reinforcement via callose deposition [50]. In this study, an enrichment analysis of DEGs under ammonium nutrition revealed that a set of overexpressed genes related to secondary cell wall materials could contribute to the maintenance of cell wall structure and the functional properties of *S. neei*. Some studies have reported that genes related to cell wall maintenance are significantly enriched in treatments that induce biotic or abiotic stress [51]. This is very interesting, as it has been shown that the cell wall structure is reassembled through the biosynthesis of cell wall polymers, which allows it to adapt to stress conditions [49, 52, 53]. Wang et al. [53] demonstrated that cell wall modifications were highly active in response to salt stress in cotton. The most common adaptations of the cell wall to stress include (i) increased levels of xyloglucan endotransglucosylase/hydrolase (XTH) and expansin proteins, associated with an increase in the degree of rhamnogalacturonan I branching, which maintains cell wall plasticity, and (ii) increased cell wall thickening through the reinforcement of the secondary wall with hemicellulose and lignin deposition [49, 54]. It was proposed that xyloglucan regulation by expansins could improve the efficiency of nutrient uptake. Several types of expansins respond to deficiencies in different nutrients, including nitrogen, phosphorus, potassium, and iron [55]. In this study, we found up-regulated genes related to the xyloglucan metabolic process, cellular–glucan metabolic process categories, which can regulate several physiological plant responses through cell wall remodeling in *S. neei*. Also, we found up-regulated genes related to sterol metabolic processes and steroid metabolic processes, which play crucial roles in various physiological and biochemical processes during development and stress resistance in plants (Fig. 4).

KEGG analysis showed that DEGs were most enriched in the biosynthesis of secondary metabolites (Table 3). This intracellular physiological defense response helps plants resist environmental stresses, indicating that they may likely play an important role in ammonium stress response, even though the shoot only receives NH_4_^+^ stress signals from the root indirectly [56]. Some studies indicate that the accumulation of secondary metabolites is highly dependent on environmental factors such as light, temperature, soil water, soil fertility, and salinity [57]. Another pathway observed in high-ammonium conditions was plant hormone signal transduction (Table 3); the abundance of DEGs related to this pathway is in agreement with the increased expression of the ethylene-responsive transcription factor and the ABA responsive element binding factor. Plant hormones play critical roles in response to various adverse biotic and abiotic environmental stresses (drought, heat, soil salinity, tropospheric ozone and excess UV radiation) [58], but have not been reported in ammonium stress. The genes involved in transcription, consistent with the role of phytohormones as triggers of signal transduction cascades, are key components in responses to abiotic stress. We investigated the common and specific responses of *S. neei* shoots to ammonium stress through up-regulated or down-regulated DEGs, with a special focus on the role of phytohormones, and we found abscisic acid (ABA) and ethylene to be essential plant hormones regulating abiotic and biotic stresses [59, 60]. According to previous studies on some halophytes species, under high-salinity conditions, ethylene production is increased in the leaves and roots [61]. Similarly, it was observed that ABA and ET signaling pathways were increased under ammonium treatment in rice and maize [23, 56], suggesting that ET and ABA biosynthesis might be increased in response to stress. It is well known that these pathways interact among themselves at various nodes, such as hormone-responsive transcription factors, to regulate plant defense responses [62-64]. Among the hormone pathways identified, ABA and ethylene biosynthesis genes were up-regulated under ammonium treatment in *S. neei*. In our study, ABA was found in one pathway (plant hormone signal transduction) and ethylene was found in three pathways (biosynthesis of secondary metabolite, plant hormone signal transduction) in the top 10 KEGG table (Table 3 and Table 4, and Supplementary Table S2). We found that S-AdoMet, SAM and ACC (direct precursor of the plant hormone ethylene) [65] (Fig. 6) are centrally involved in ethylene biosynthesis, and ZEP, VDE1 and ABA2 are involved in ABA biosynthesis. In addition, we found the key ABA responsive element binding factor and ethylene-responsive transcription factor. Therefore, it seems that the transduction pathway, regulated by ABA and ethylene, plays an important role in the stress response of *S*.*neei* to NH_4_^+^.

**Table 4.**
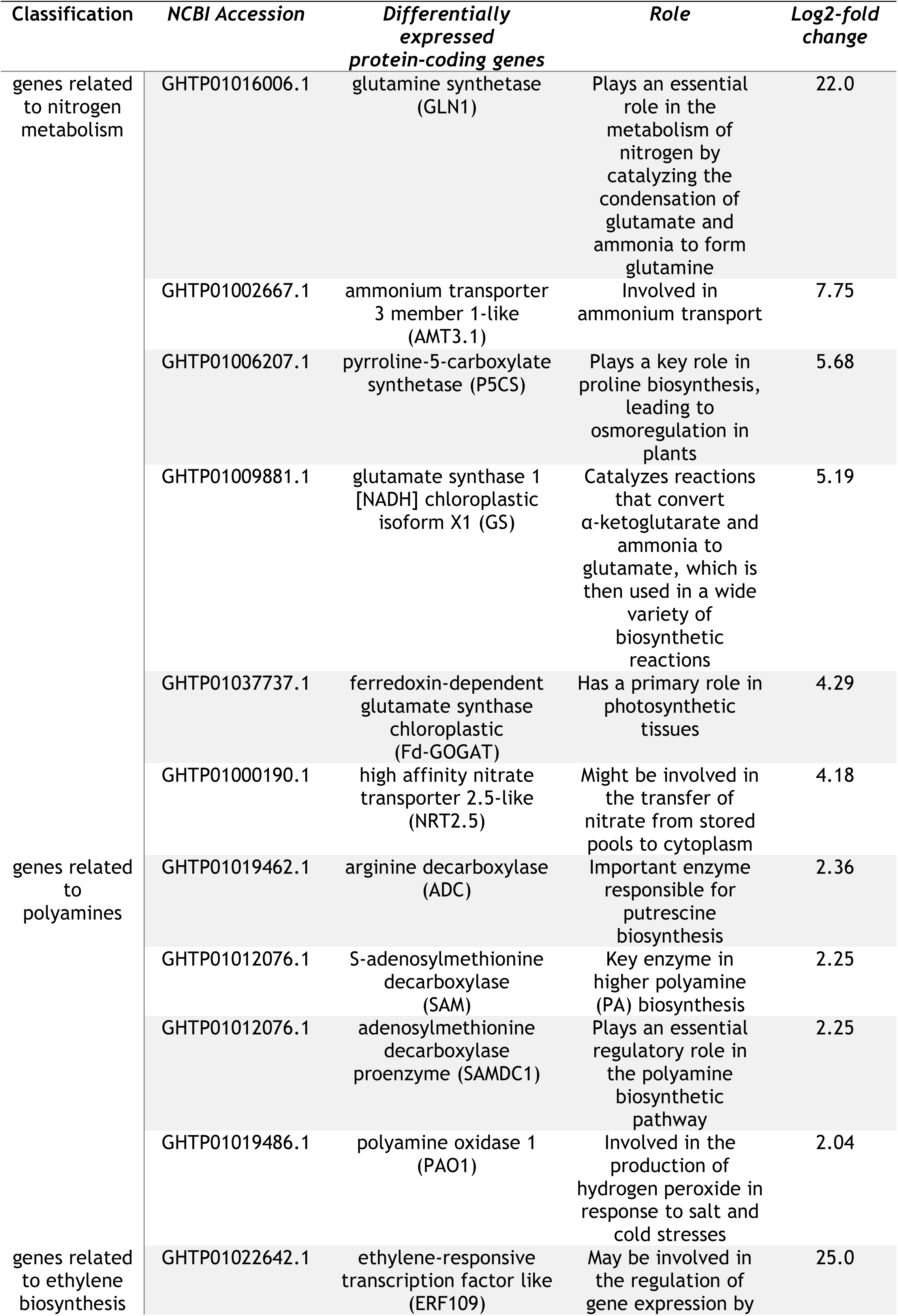
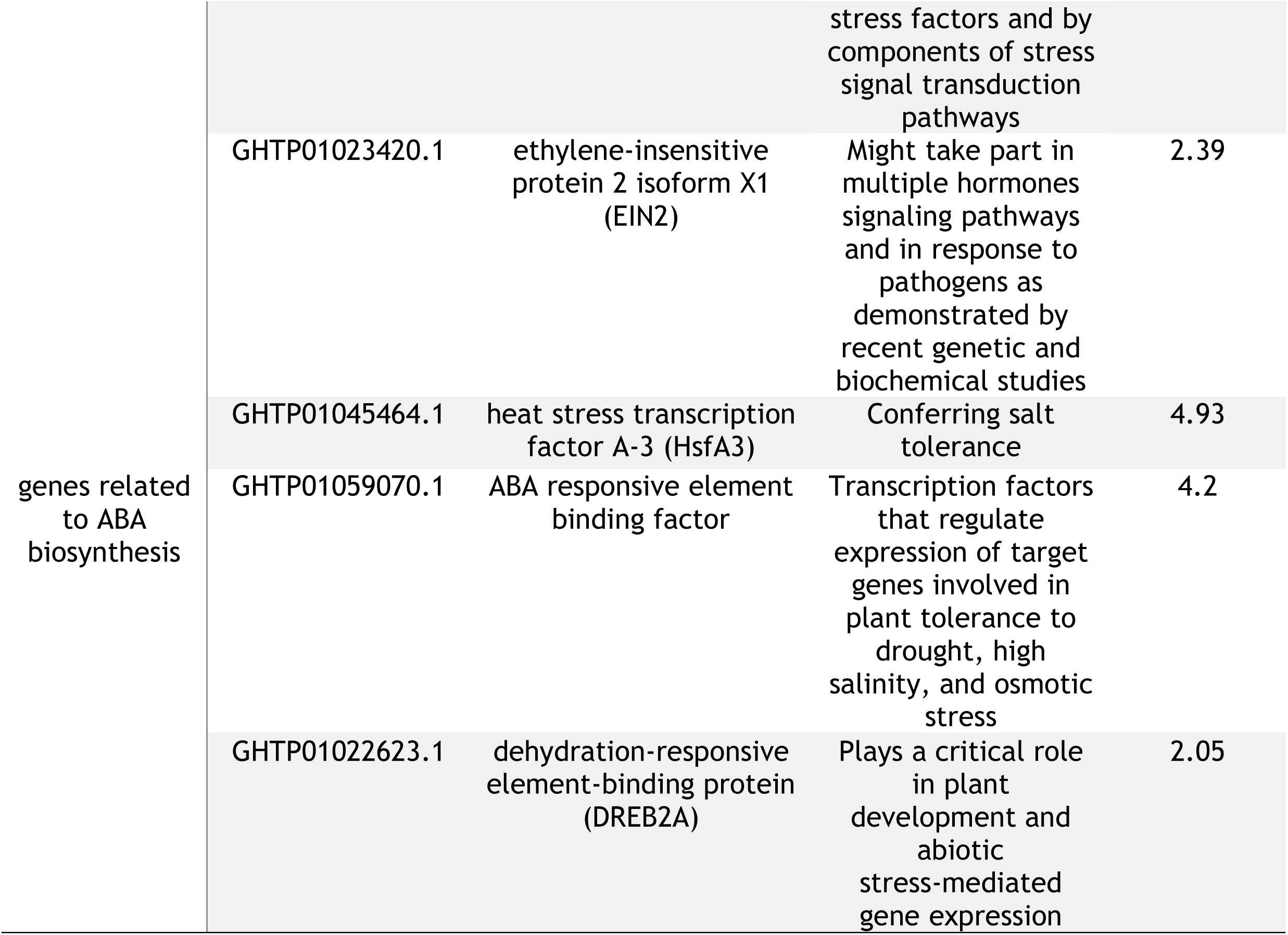
Genes involved in ammonium homeostasis and phytohormone biosynthesis in *Salicornia neei*.

**Fig. 6.**
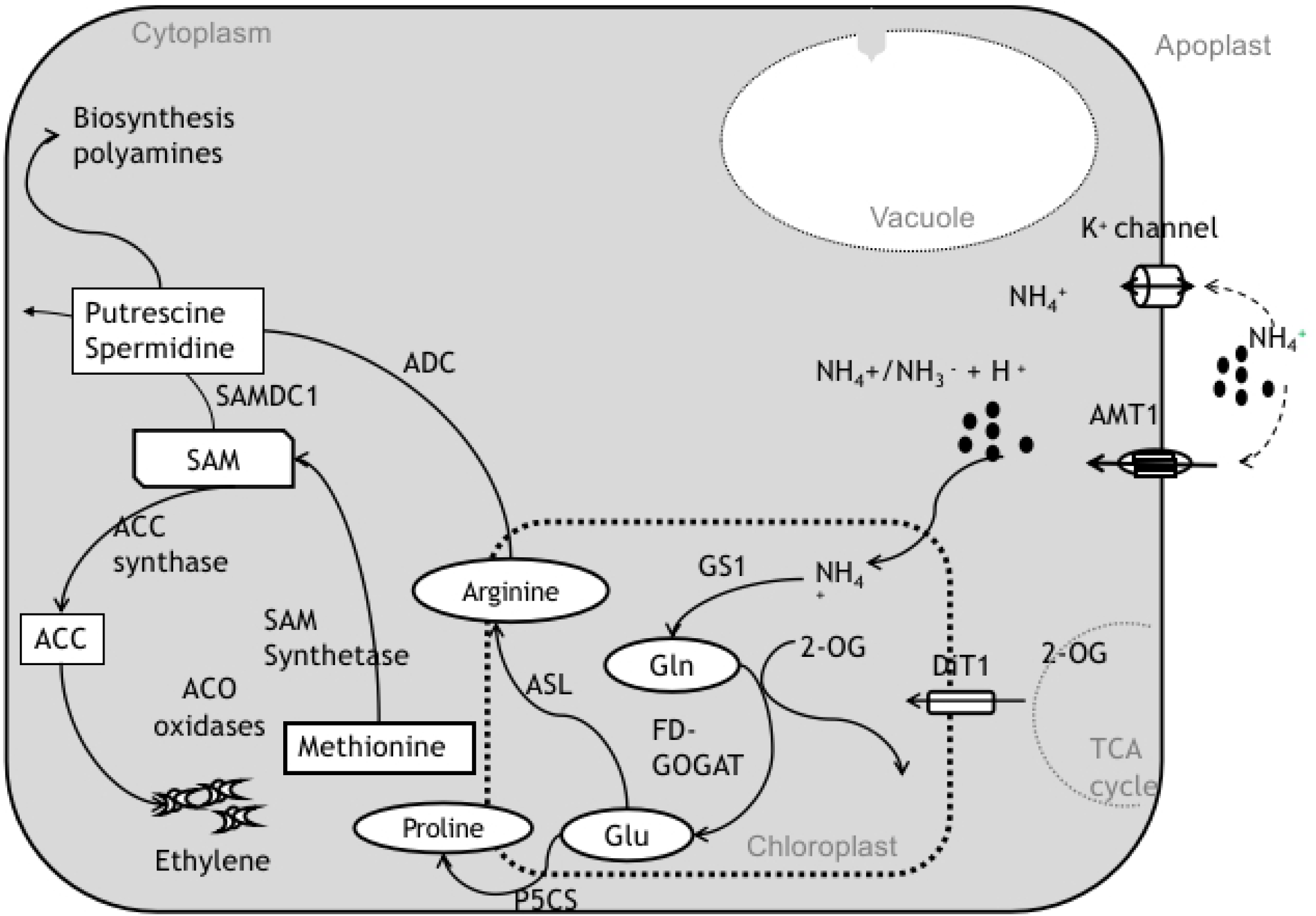
Putative model of ammonium metabolism in *Salicornia neei*, polyamines and ethylene biosynthesis, according to the major DEGs that were modulated by NH4+: ammonium transporter 1 (AMT1), glutamine synthetase 1 (GS1), ferredoxin-dependent glutamate synthase (Fd-GOGAT), glutamate (Glu), glutamine (Gln), pyrroline-5-carboxylate synthetase (P5CS), 2-oxoglutarate (2-OG), dicarboxylate transporter 1 (DiT1), arginine decarboxylase (ADC), S-adenosylmethionine decarboxylase (SAMDC1), tricarboxylic acid cycle (TCA), and potassium K^+^ channel, ACO, ACC. Figure adapted from Ma et al. [30] and Chen et al [60].

The biosynthesis pathways of polyamines (PAs) and ethylene are interrelated, with SAM as a common precursor that can be used to form ACC, the precursor of ethylene that is active in the conversion of PAs [66]. Their physiological functions are distinct and at times antagonistic [67], but both have been identified as important signaling molecules involved in stress tolerance [68]. In this study, overexpressed genes related to the metabolic pathways of polyamines were found (Table 4, Fig. 6, and Supplementary Table S2). According to Navin et al. [69], putrescine, spermidine, and spermine biosynthesis are also involved in the amelioration of drought, salinity, cold, and heat stresses. Polyamines counter stress by binding with nucleic acids, proteins, and phospholipids to stabilize their structures in response to diverse abiotic stress conditions [69]. Bouchereau et al. [70] stated that NH_4_ ^+^ nutrition is associated with significant changes in the free polyamine content in the shoots or roots of plants, which could be key to the protection of a stressed cell. We found three genes, PAO1, PAO2 and PAO4, encoding polyamine oxidase (PAO), which is an enzyme with distinct physiological roles that is responsible for polyamine catabolism. In plants, increasing evidence suggests that PAO genes play essential roles in the abiotic and biotic stress response, as some PAOs catalyze the reverse reaction of PA synthesis via the PA back-conversion pathway [71]. Tavladoraki et al. [72] identified PAO1 in Arabidopsis as an enzyme that possesses back-conversion capacity, responsible for the conversion of Spm to Spd. Similarly, Moschou el al. [73] identified that, in Arabidopsis, PAO1 and PAO4 were able to convert Spm to Spd, and PAO2 and PAO3 catalyzed the production of Spd from Spm before producing Put. This result suggests the involvement of PAO genes in stress responses, and their probable implications for PA homeostasis in *S. neei*.

Considering the possible abiotic stress induced by ammonium-based nutrition, we looked for processes activated in the plant that help to avoid ammonium accumulation or to mitigate its effects. Thus, sets of 72 up-regulated and 19 down-regulated genes related to nitrogen metabolism were found to be modulated by NH_4_^+^ at salinity concentrations close to 600 mM NaCl in the shoots. Among the main genes recognized in the metabolism of NH_4_^+^ in plants, we propose a possible route based on several up-regulated genes, such as GS, GLT-1, 2-oxoglutarate dehydrogenase (2-OGD), ferredoxin-dependent glutamate synthase chloroplastic (Fd-GOGAT), and glutamate synthase 1 (NADH) chloroplastic isoform X1 (GLT1) (see Table 4, Fig. 6 and Supplementary Table S2). Similar results have been published concerning the halophyte plant *Salicornia europeae*, which showed high GS and GDH activity at high salinity concentrations [30]. Further, Ma et al. [30] noted that *Salicornia* plants fed with ammonia also require salinity of 200 mM NaCl or more in the substrate, conditions that can presumably stimulate the detoxification mechanisms generated by stress [74]. This could explain the ability of halophyte plants to thrive in environments where salinity is high and ammonia is a common nutrient. It has previously been revealed that some wetland plants and other marine species that grow in terrestrial habitats where NH_4_ ^+^ prevails over NO_3_ ^−^ have a special preference for ammonia [75].

In plants, diverse abiotic stresses have been shown to induce autophagy, including nutrient starvation and oxidative stress, improving plant resistance [76]. The energy sensors SNF-related kinase 1 (SnRK1) and target of rapamycin (TOR) control autophagy under energy deficiency, but also under diverse stress conditions [77]. Signorelli et al. [77] have suggested that the accumulation of GABA, PA, ethylene and ABA under stress conditions can indirectly control autophagy as well, by different pathways.

Studies have theorized that additional cytosolic nutrients can be maintained by compartmentalizing them at sites where they are not metabolized, such as the vacuole [78, 79]. This direct encapsulation of nitrogen is possible through specialized autophagic vesicles that subsequently fuse with the vacuole for proteolysis and hydrolysis [80]. Moreover, materials such as nutrients can be remobilized to new growing organs and sinks, such as seeds. In Arabidopsis, it was observed that autophagy can regulate the abiotic stress caused by the excessive uptake of toxic ions; researchers reported that the level of autophagy peaks within 30 min after salt stress, and then settles into a new homeostasis, but such an induction is absent in defective mutants in autophagy [81]. In this transcriptome, we found a series of autophagy-related genes (ATGs) that are responsible for the initiation and formation of autophagosomes, such as the following: the ATG1/ATG13 kinase complex that initiates autophagosome formation; 2) ATG9 forms a complex with accessary proteins ATG2 and ATG18, which promotes phagophore expansion and has been described as the only integral complex necessary for the formation of the autophagosomal membrane [82]; 3) a complex for vesicle nucleation; 4) the ATG8/PE conjugates, which are located in both the inner and the outer autophagosome membranes, aid phagophore expansion and vesicle closure to form autophagosomes, and recognize autophagic cargoes through ATG8-interacting proteins [83]. These genes related to autophagy are included in the autophagy pathway and others (Table 3). This vacuolar complex is probably induced to avoid apoplastic acidification, which generates stress and leaf senescence [28, 84, 85] (see Fig. 7).

**Fig. 7.**
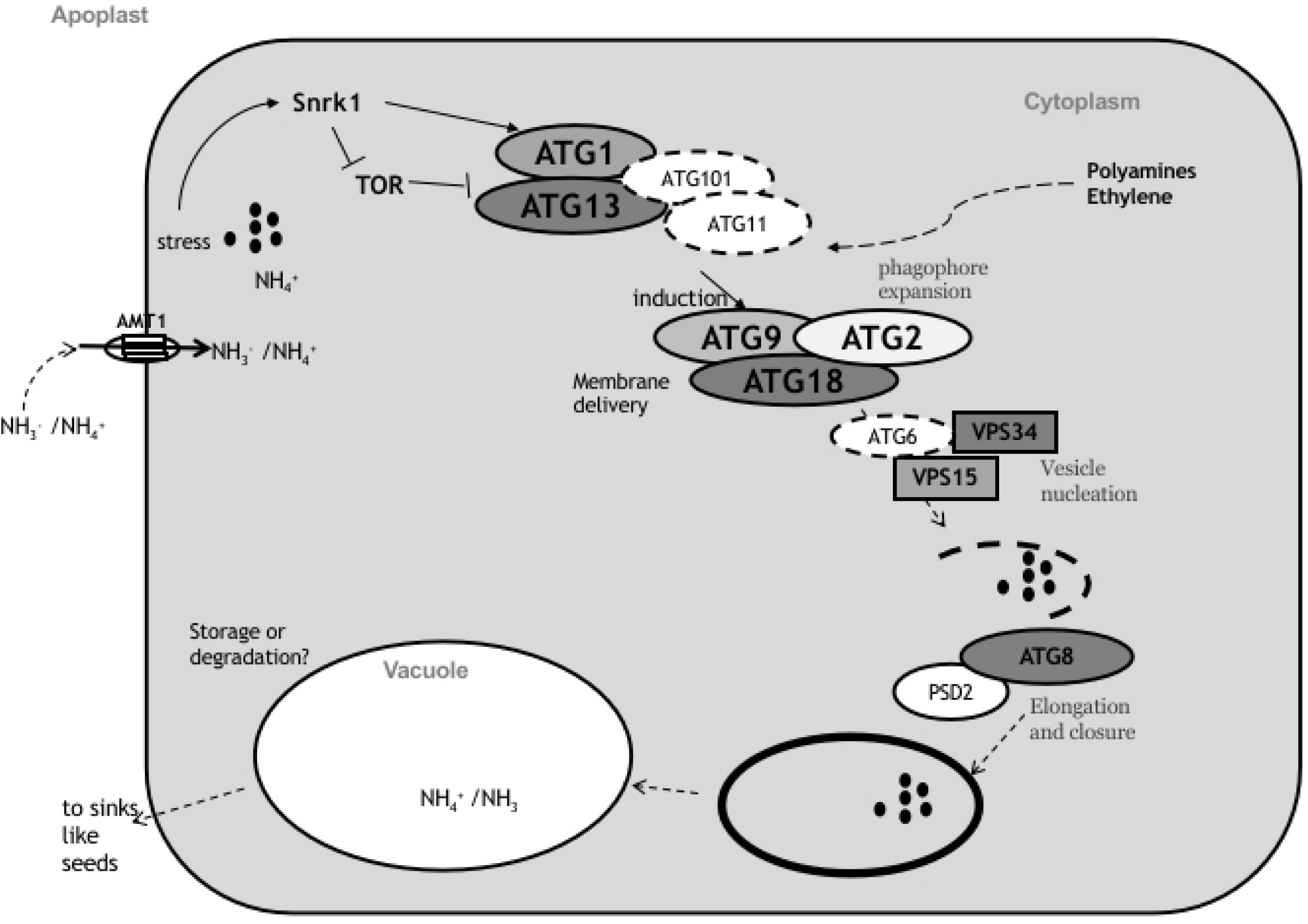
Putative schematic diagram of the autophagy process in *Salicornia neei*. Energy sensors SNF-related kinase 1 (SnRK1), target of rapamycin (TOR), autophagy-related protein 1 (ATG1), autophagy-related protein 11 (ATG11), autophagy-related protein 101 (ATG101), autophagy-related protein 13 (ATG13), autophagy-related protein 9 (ATG9), autophagy-related protein 2 (ATG2), autophagy-related protein 18, vacuolar protein sorting 34 (VPS34), vacuolar protein sorting 15 (VPS15), autophagy-related protein 6 (ATG6), protein sorting 8 (ATG8), phosphatidylserine decarboxylase proenzyme 2 (PSD2) (2.0-fold). In dotted oval ATG not observed. Figure adapted from Chen et al.[80].

## 5. Conclusions

The results suggest that the ammonium detoxification system in *S. neei* is mediated by a wide variety of up-regulated genes that are associated with the maintenance of ammonium homeostasis through the activation of glutamine and glutamate synthetase, accompanied by the biosynthesis of the phytohormones and polyamines involved in the protection of important protein structures under stress conditions.

In *S. neei*, the vacuolar complex was probably created to avoid the apoplastic acidification induced by ammonium nutrition.

Cell wall biosynthesis may correspond to the stress response induced by excess ammonia, taking into account the fact that several studies addressing stress response in plants have reported the biosynthesis of cell wall structures.

The kinetics of ammonium uptake by *Salicornia neei* under ammonium nutrition have been characterized using the Michaelis–Menten equation. Our results support the hypothesis that *S. neei* is very effective in removing ammonium at concentrations < 4mM, compared to other species used in aquaculture wastewater [7, 86]. The maximum rates (Imax) and the half-saturation constants (Km) for ammonium uptake were identified, demonstrating that *S. neei* can be used to treat effluents with ammonium pollutants. *Salicornia neei* showed a high affinity for the substrate, which can be used to decontaminate waters with high nutrient loads derived from marine aquaculture.

## Acknowledgments

The authors wish to thank Aldo Madrid and Marine Farms Inc. for their technical assistance and support during the project. M.R.D. was supported by a Doctoral fellowship from the “Dirección General de Investigación y Postgrado” of the Universidad Técnica Federico Santa María.

