## Supplementary material for "RNA-Seq analysis and transcriptome assembly of *Salicornia neei* reveals a powerful system for ammonium detoxification": supplementarymaterial.docx

**Table S1.** Generalized linear model for analysis of ammonium removal rate; the results reveal that the ammonium removal rate is significantly different from 0 NH_4_Cl for concentrations of 1, 2, 3, and 4 mM ammonium.

| **Concentration**  **NH_4_ (mM)** | **Ammonium removal rate**  **(µmol min^−1^)** | **SD** | **t value** | **p (> t\|)** |
| --- | --- | --- | --- | --- |
| 0 | −0.01085 | 0.01253 | −0.866 | 0.407 |
| 1 | −0.36194 | 0.04676 | −7.741 | 1.57e−05 |
| 2 | −0.4757 | 0.1247 | −3.814 | 0.00341 |
| 3 | −0.5136 | 0.1332 | −3.856 | 0.00318 |
| 4 | −0.5602 | 0.1260 | −4.445 | 0.00124 |

SD: standard deviation; t value: t-statistic; p: probability.

**Table S2**. Genes related to nitrogen metabolism and phytohormone biosynthesis.

| ID_NCBI | Description | Fold change | FDR -p value |
| --- | --- | --- | --- |
| GHTP01000190.1 | high affinity nitrate transporter 2.5-like | 4.18 | 0.00E+00 |
| GHTP01000614.1 | RNI-like superfamily protein | 2.72 | 0 |
| GHTP01001242.1 | probable polyamine oxidase 2 | 2.42 | 0 |
| GHTP01001331.1 | ABSCISIC ACID-INSENSITIVE 5-like protein 2 | -2.01 | 9.13E-08 |
| GHTP01001870.1 | ethylene-responsive transcription factor 2-like | 2.06 | 0.00E+00 |
| GHTP01001939.1 | annexin D2-like | -2.16 | 0.00E+00 |
| GHTP01002359.1 | arginine/serine-rich coiled-coil protein 2-like isoform X2 | -2.64 | 0.00E+00 |
| GHTP01002427.1 | probable protein phosphatase 2C 25 | 4.29 | 4.12E-06 |
| GHTP01002474.1 | ribosome biogenesis protein NSA2 homolog | -2.66 | 1.91E-14 |
| GHTP01002667.1 | ammonium transporter 3 member 1-like | 7.75 | 3.22E-04 |
| GHTP01002690.1 | probable 2-oxoglutarate-dependent dioxygenase At5g05600 | 2.42 | 0.00E+00 |
| GHTP01002692.1 | ferredoxin--nitrite reductase. chloroplastic | 2.85 | 0.00E+00 |
| GHTP01002763.1 | plastidal glycolate/glycerate translocator 1. chloroplastic | 7.45 | 0.00E+00 |
| GHTP01002899.1 | probable polyamine oxidase 2 | 3.04 | 0 |
| GHTP01002922.1 | hypothetical protein C5167_007012 | -2.67 | 3.11E-06 |
| GHTP01003383.1 | lipase-like PAD4 | 2.02 | 1.89E-09 |
| GHTP01003587.1 | Ninja-family protein AFP3 | -2.03 | 6.84E-04 |
| GHTP01003789.1 | pathogenesis-related protein STH-21-like | 2.47 | 1.63E-12 |
| GHTP01003798.1 | ethylene-overproduction protein 1 | 2.3 | 9.18E-07 |
| GHTP01003959.1 | protein NRT1/ PTR FAMILY 6.1-like | 3.39 | 2.15E-06 |
| GHTP01004213.1 | protein NRT1/ PTR FAMILY 6.3 | 3.14 | 0.00E+00 |
| GHTP01004746.1 | 1-aminocyclopropane-1-carboxylate synthase | 25.0 | 2.90E-06 |
| GHTP01004887.1 | glutamate synthase 1 [NADH]. chloroplastic isoform X1 | 6.74 | 2.61E-06 |
| GHTP01004888.1 | glutamate synthase 1 [NADH]. chloroplastic isoform X1 | 3.68 | 1.87E-05 |
| GHTP01005340.1 | calcineurin-like metallo-phosphoesterase superfamily protein | 3.24 | 0 |
| GHTP01005543.1 | arginine/serine-rich coiled-coil protein 2-like isoform X2 | -3.19 | 0.00E+00 |
| GHTP01005852.1 | probable ethylene response sensor 1 | -2.58 | 2.90E-06 |
| GHTP01005883.1 | ferredoxin--NADP reductase. leaf-type isozyme. chloroplastic | 5.8 | 0.00E+00 |
| GHTP01006207.1 | delta-1-pyrroline-5-carboxylate synthase | 5.68 | 0.00E+00 |
| GHTP01006464.1 | Protein kinase superfamily protein with octicosapeptide/Phox/Bem1p domain | 4.04 | 0.00021 |
| GHTP01006862.1 | probable serine/threonine-protein kinase At4g35230 | -2.1 | 3.03E-06 |
| GHTP01006991.1 | high affinity nitrate transporter 2.5-like | 2.92 | 4.59E-08 |
| GHTP01007207.1 | Alpha-D-phosphohexomutase superfamily | 14.48 | 0 |
| GHTP01007773.1 | protein NRT1/ PTR FAMILY 5.10-like | 2.5 | 1.62E-13 |
| GHTP01007774.1 | protein NRT1/ PTR FAMILY 3.1-like | 2.16 | 0.00E+00 |
| GHTP01007786.1 | sulfite reductase 1 [ferredoxin]. chloroplastic | 2.96 | 7.28E-13 |
| GHTP01007787.1 | sulfite reductase 1 [ferredoxin]. chloroplastic | 2.77 | 2.71E-14 |
| GHTP01008037.1 | probable histone-arginine methyltransferase 1.3 | 2.81 | 0.00E+00 |
| GHTP01008147.1 | protein kinase superfamily protein | 2.25 | 0.00000104 |
| GHTP01008259.1 | 1-aminocyclopropane-1-carboxylate synthase | 13.96 | 0.00E+00 |
| GHTP01008429.1 | systemin receptor SR160 | 4.09 | 0.00E+00 |
| GHTP01008430.1 | systemin receptor SR160-like | 3.79 | 1.50E-10 |
| GHTP01008433.1 | leucine-rich repeat receptor-like serine/threonine-protein kinase BAM1 | 3.15 | 6.47E-05 |
| GHTP01008449.1 | piriformospora indica-insensitive protein 2 | -3.45 | 5.86E-11 |
| GHTP01008618.1 | 2-oxoglutarate dehydrogenase. mitochondrial-like | 2.34 | 0.00E+00 |
| GHTP01009131.1 | protein NRT1/ PTR FAMILY 6.4 | 4.23 | 1.15E-07 |
| GHTP01009152.1 | threonine dehydratase biosynthetic. chloroplastic-like | 2.61 | 1.18E-06 |
| GHTP01009640.1 | 24-methylenesterol C-methyltransferase 2 | 3.79 | 0.00E+00 |
| GHTP01009702.1 | putative glutamine amidotransferase GAT1_2.1 | 2.47 | 6.33E-06 |
| GHTP01009881.1 | glutamate synthase 1 [NADH]. chloroplastic isoform X1 | 5.19 | 0.00E+00 |
| GHTP01009884.1 | glutamine--fructose-6-phosphate aminotransferase [isomerizing] 2 | 2.27 | 2.45E-04 |
| GHTP01010361.1 | DEAD-box ATP-dependent RNA helicase 38 | 2.63 | 0.00E+00 |
| GHTP01010892.1 | phenylalanine ammonia-lyase | 2.56 | 0.00E+00 |
| GHTP01010893.1 | phenylalanine ammonia-lyase | 2.7 | 0.00E+00 |
| GHTP01010894.1 | phenylalanine ammonia-lyase | 2.22 | 0.00E+00 |
| GHTP01010915.1 | ammonium transporter 1 member 1 | 2.39 | 0.00E+00 |
| GHTP01011282.1 | S-adenosylmethionine synthase 1 | 2.22 | 0.000000618 |
| GHTP01012076.1 | S-adenosylmethionine decarboxylase proenzyme | 2.25 | 0 |
| GHTP01012190.1 | asparagine synthetase [glutamine-hydrolyzing] 2 | 2.39 | 3.50E-10 |
| GHTP01012332.1 | aldehyde dehydrogenase family 3 member H1-like isoform X1 | 2.98 | 0.00E+00 |
| GHTP01012347.1 | delta-1-pyrroline-5-carboxylate synthase isoform X1 | 2.1 | 0.00E+00 |
| GHTP01012348.1 | delta-1-pyrroline-5-carboxylate synthase isoform X1 | 3.63 | 2.91E-13 |
| GHTP01012357.1 | delta-1-pyrroline-5-carboxylate synthase isoform X2 | 2.41 | 0.00E+00 |
| GHTP01012664.1 | protein TRANSPARENT TESTA GLABRA 1-like | 2.24 | 3.03E-09 |
| GHTP01012984.1 | glycosyltransferase family 64 protein C4 | 2.27 | 0.00E+00 |
| GHTP01013071.1 | putative glutamine synthetase | 18.0 | 9.11E-05 |
| GHTP01013078.1 | chloride channel protein CLC-b | 4.28 | 1.69E-06 |
| GHTP01013407.1 | glutamate synthase 1 [NADH]. chloroplastic isoform X1 | 2.14 | 1.46E-08 |
| GHTP01013581.1 | 24-methylenesterol C-methyltransferase 2 | 3.03 | 0.00E+00 |
| GHTP01013791.1 | receptor-like protein kinase HSL1 | 4.04 | 9.90E-07 |
| GHTP01013796.1 | probable leucine-rich repeat receptor-like protein kinase At5g63930 | 4.72 | 5.56E-04 |
| GHTP01014868.1 | serine/threonine-protein kinase TOR | 2.73 | 1.36E-11 |
| GHTP01015115.1 | DEAD-box ATP-dependent RNA helicase 38 | 2.02 | 9.04E-11 |
| GHTP01015744.1 | beta-amyrin synthase | 2.55 | 2.33E-06 |
| GHTP01015954.1 | nitrate reductase | 2.01 | 3.90E-10 |
| GHTP01016006.1 | glutamine synthetase | 22.0 | 1.27E-05 |
| GHTP01016118.1 | interferon-related developmental regulator 1 | 2.64 | 0 |
| GHTP01016249.1 | plastidal glycolate/glycerate translocator 1. chloroplastic | 4.01 | 0.00E+00 |
| GHTP01016476.1 | ferredoxin-dependent glutamate synthase. chloroplastic | 3.07 | 0.00E+00 |
| GHTP01017187.1 | Alpha-D-phosphohexomutase superfamily | 4.43 | 0 |
| GHTP01017631.1 | probable 2-oxoglutarate-dependent dioxygenase At5g05600 | 2.04 | 0.00E+00 |
| GHTP01017820.1 | Calcineurin-like metallo-phosphoesterase superfamily protein | 2.67 | 1.99E-09 |
| GHTP01018073.1 | formamidase-like isoform X2 | 2.16 | 2.50E-07 |
| GHTP01018144.1 | heptahelical transmembrane protein 2-like | -2.74 | 0.00E+00 |
| GHTP01018148.1 | heptahelical transmembrane protein 4-like isoform X1 | 2.05 | 0.00E+00 |
| GHTP01018149.1 | heptahelical transmembrane protein 1 | 2.92 | 2.85E-06 |
| GHTP01018169.1 | transmembrane 9 superfamily member 3 | 3.23 | 0 |
| GHTP01018170.1 | transmembrane 9 superfamily member 3 | 5.36 | 0 |
| GHTP01018171.1 | transmembrane 9 superfamily member 3 | 5.62 | 0 |
| GHTP01018185.1 | transmembrane 9 superfamily member 3 | 2.04 | 0 |
| GHTP01018187.1 | transmembrane 9 superfamily member 8 | 3.16 | 1.91E-14 |
| GHTP01018643.1 | transcription termination factor MTERF2. chloroplastic | 2.24 | 0.000000242 |
| GHTP01019001.1 | abscisic stress ripening | 2.35 | 0.00E+00 |
| GHTP01019273.1 | 24-methylenesterol C-methyltransferase 2 | 2.75 | 0.00E+00 |
| GHTP01019434.1 | histone deacetylase 6 | 3.77 | 5.10E-06 |
| GHTP01019462.1 | arginine decarboxylase | 2.36 | 0.00E+00 |
| GHTP01019486.1 | polyamine oxidase 1 | 2.04 | 0.0000346 |
| GHTP01019489.1 | probable polyamine oxidase 2 | 4.6 | 0 |
| GHTP01019491.1 | probable polyamine oxidase 2 | 3.3 | 1.27E-11 |
| GHTP01020110.1 | glutamate synthase 1 [NADH]. chloroplastic-like isoform X3 | 2.23 | 0.00E+00 |
| GHTP01020228.1 | abscisic acid 8'-hydroxylase 1-like | 3.03 | 5.07E-05 |
| GHTP01020513.1 | probable leucine-rich repeat receptor-like protein kinase At2g33170 | 2.82 | 1.93E-10 |
| GHTP01020930.1 | E3 ubiquitin-protein ligase XBAT32-like | 2.23 | 1.09E-04 |
| GHTP01021097.1 | Urease | 4.76 | 1.49E-04 |
| GHTP01021384.1 | threonine dehydratase biosynthetic. chloroplastic-like | 4.2 | 0.00E+00 |
| GHTP01021416.1 | ammonium transporter 3 member 1-like | -2.06 | 0.00E+00 |
| GHTP01021421.1 | dicarboxylate transporter 2.1. chloroplastic-like | 2.24 | 0.00E+00 |
| GHTP01021422.1 | dicarboxylate transporter 1. chloroplastic | 2.21 | 0.00E+00 |
| GHTP01021423.1 | dicarboxylate transporter 2.1. chloroplastic-like | 2.87 | 0.00E+00 |
| GHTP01021425.1 | dicarboxylate transporter 1. chloroplastic | 2.35 | 0.00E+00 |
| GHTP01021568.1 | E3 ubiquitin-protein ligase BAH1-like | 2.13 | 6.18E-13 |
| GHTP01021801.1 | histidine kinase 2 | -2.09 | 6.38E-04 |
| GHTP01022325.1 | probable polyamine oxidase 4 | 6.98 | 0 |
| GHTP01022354.1 | receptor-like protein kinase HSL1 | 3.41 | 2.29E-05 |
| GHTP01022355.1 | probable leucine-rich repeat receptor-like protein kinase At5g63930 | 3.45 | 9.46E-06 |
| GHTP01022604.1 | ethylene-responsive transcription factor ERF003-like | 2.28 | 7.33E-06 |
| GHTP01022606.1 | ethylene-responsive transcription factor ERF054-like | 5.13 | 0.00E+00 |
| GHTP01022607.1 | ethylene-responsive transcription factor CRF2-like | 2.51 | 1.47E-06 |
| GHTP01022608.1 | AP2-like ethylene-responsive transcription factor ANT | -3.16 | 1.45E-15 |
| GHTP01022611.1 | ethylene-responsive transcription factor RAP2-7-like isoform X4 | 2.65 | 0.00E+00 |
| GHTP01022615.1 | ethylene-responsive transcription factor TINY-like | 6.4 | 0.00E+00 |
| GHTP01022617.1 | Ethylene-responsive transcription factor RAP2-10 | 2.8 | 0.00E+00 |
| GHTP01022622.1 | ethylene-responsive transcription factor RAP2-4-like | 3.16 | 0.00E+00 |
| GHTP01022624.1 | ethylene-responsive transcription factor ERF113-like | 2.36 | 0.00E+00 |
| GHTP01022626.1 | ethylene-responsive transcription factor ERF113 | 2.09 | 0.00E+00 |
| GHTP01022627.1 | ethylene-responsive transcription factor ERF071 | 2.97 | 0.00E+00 |
| GHTP01022630.1 | AP2-like ethylene-responsive transcription factor At1g16060 | 2.12 | 2.18E-05 |
| GHTP01022635.1 | AP2-like ethylene-responsive transcription factor AIL6 | 2.45 | 2.95E-04 |
| GHTP01022637.1 | ethylene-responsive transcription factor TINY-like | 12.13 | 6.62E-05 |
| GHTP01022638.1 | ethylene-responsive transcription factor ERF017 | 22.23 | 2.74E-13 |
| GHTP01022640.1 | ethylene-responsive transcription factor 5 | 2.88 | 0.00E+00 |
| GHTP01022641.1 | ethylene-responsive transcription factor 4 | 10.05 | 0.00E+00 |
| GHTP01022642.1 | ethylene-responsive transcription factor ERF109-like | 25.0 | 2.90E-06 |
| GHTP01022644.1 | ethylene-responsive transcription factor RAP2-4-like | 2.65 | 0.00E+00 |
| GHTP01022648.1 | ethylene-responsive transcription factor ERF114 | 3.97 | 0.00E+00 |
| GHTP01022649.1 | ethylene-responsive transcription factor ERF114 | 3.93 | 1.11E-14 |
| GHTP01022650.1 | ethylene-responsive transcription factor ERF073-like | 3.96 | 0.00E+00 |
| GHTP01022652.1 | ethylene-responsive transcription factor 1B-like | 4.31 | 0.00E+00 |
| GHTP01023041.1 | leucine-rich repeat receptor-like serine/threonine-protein kinase BAM1 | -2.14 | 0.00E+00 |
| GHTP01023305.1 | abscisic acid 8'-hydroxylase 1-like | 2.65 | 2.18E-14 |
| GHTP01023397.1 | pyrroline-5-carboxylate synthetase | 3.08 | 0.00E+00 |
| GHTP01023406.1 | acetylglutamate kinase. chloroplastic | -4.67 | 1.96E-04 |
| GHTP01023416.1 | delta-1-pyrroline-5-carboxylate synthase isoform X2 | 3.3 | 0.00E+00 |
| GHTP01023420.1 | ethylene-insensitive protein 2 isoform X1 | 2.39 | 0.00E+00 |
| GHTP01023431.1 | ethylene-insensitive protein 2 | -2.05 | 0.00E+00 |
| GHTP01023877.1 | CBL-interacting protein kinase 2-like | 2.48 | 4.71E-10 |
| GHTP01023878.1 | CBL-interacting protein kinase 2-like | 2.83 | 3.68E-10 |
| GHTP01023927.1 | leucine-rich repeat receptor-like serine/threonine-protein kinase BAM1 | -2.14 | 8.06E-07 |
| GHTP01023947.1 | protein kinase-like | 2.32 | 1.50E-06 |
| GHTP01024045.1 | bifunctional protein FolD 4. chloroplastic-like isoform X2 | 2.06 | 4.65E-09 |
| GHTP01024949.1 | protein NRT1/ PTR FAMILY 1.1-like | 3.52 | 0.00E+00 |
| GHTP01024954.1 | protein NRT1/ PTR FAMILY 3.1 | 2.89 | 0.00E+00 |
| GHTP01024957.1 | protein NRT1/ PTR FAMILY 6.3 | 2.04 | 7.74E-06 |
| GHTP01024969.1 | protein NRT1/ PTR FAMILY 1.2-like | 2.79 | 2.87E-15 |
| GHTP01024973.1 | protein NRT1/ PTR FAMILY 3.1-like | 2.45 | 3.81E-13 |
| GHTP01024977.1 | protein NRT1/ PTR FAMILY 1.1-like | 19.2 | 7.37E-08 |
| GHTP01024978.1 | protein NRT1/ PTR FAMILY 1.1-like | 6.28 | 2.71E-12 |
| GHTP01025534.1 | threonine dehydratase biosynthetic. chloroplastic-like | 4.17 | 0.00E+00 |
| GHTP01025626.1 | cytochrome P450 78A7-like | 4.85 | 2.87E-06 |
| GHTP01025695.1 | PAP/OAS1 substrate-binding domain superfamily | 2.02 | 1.22E-08 |
| GHTP01025875.1 | mitogen-activated protein kinase kinase 3 | 2.01 | 0.00E+00 |
| GHTP01026039.1 | probable histone-arginine methyltransferase 1.3 | 2.59 | 1.90E-10 |
| GHTP01026058.1 | Molecular chaperone (DnaJ superfamily) | 2.05 | 0.00000418 |
| GHTP01026231.1 | EID1-like F-box protein 3 | 2.01 | 2.87E-15 |
| GHTP01026264.1 | high affinity nitrate transporter 2.5-like | 2.28 | 0.00E+00 |
| GHTP01026297.1 | protein NRT1/ PTR FAMILY 6.1-like | 2.26 | 8.06E-05 |
| GHTP01026298.1 | protein NRT1/ PTR FAMILY 6.4 | 2.07 | 0.00E+00 |
| GHTP01026302.1 | protein NRT1/ PTR FAMILY 1.1-like | 5.84 | 0.00E+00 |
| GHTP01026539.1 | dual specificity protein kinase YAK1 homolog | 2.52 | 4.04E-07 |
| GHTP01026808.1 | ETHYLENE INSENSITIVE 3-like 1 protein | 2.21 | 0.00E+00 |
| GHTP01026846.1 | glutamate synthase 1 [NADH]. chloroplastic isoform X1 | 2.42 | 0.00E+00 |
| GHTP01026872.1 | 1-aminocyclopropane-1-carboxylate oxidase | 5.4 | 0.00E+00 |
| GHTP01027339.1 | interferon-related developmental regulator 1 | 2.42 | 0 |
| GHTP01027367.1 | ferredoxin-dependent glutamate synthase. chloroplastic | 2.14 | 0.00E+00 |
| GHTP01027484.1 | ABSCISIC ACID-INSENSITIVE 5-like protein 2 | -2.1 | 2.22E-04 |
| GHTP01027496.1 | ABSCISIC ACID-INSENSITIVE 5-like protein 5 | -3.44 | 0.00E+00 |
| GHTP01027510.1 | RNA polymerase II C-terminal domain phosphatase-like 1 | 2.65 | 0.00E+00 |
| GHTP01027622.1 | polyamine oxidase 1 | 3.6 | 0 |
| GHTP01027790.1 | threonine dehydratase biosynthetic. chloroplastic-like | 2.56 | 0.00E+00 |
| GHTP01027841.1 | ethylene-responsive transcription factor ERF054-like | 11.45 | 1.76E-06 |
| GHTP01027843.1 | ethylene-responsive transcription factor 1B-like | 4.77 | 0.00E+00 |
| GHTP01027844.1 | ethylene-responsive transcription factor ERF105-like | 3.66 | 0.00E+00 |
| GHTP01028224.1 | ABSCISIC ACID-INSENSITIVE 5-like protein 5 | -2.02 | 0.00E+00 |
| GHTP01028896.1 | polyamine oxidase-like | 17.56 | 0 |
| GHTP01029071.1 | protein NRT1/ PTR FAMILY 3.1 | 2.1 | 0.00E+00 |
| GHTP01029578.1 | nodulin-related protein 1-like | 2.51 | 0.00E+00 |
| GHTP01030137.1 | protein FAF-like. chloroplastic | 2.2 | 5.62E-15 |
| GHTP01030179.1 | nitrate regulatory gene2 protein-like | -2.28 | 0.00E+00 |
| GHTP01030608.1 | high-affinity nitrate transporter-activating protein 2.1-like | 3.12 | 0.00E+00 |
| GHTP01030932.1 | RING/FYVE/PHD zinc finger superfamily protein isoform 1 | 4.36 | 0 |
| GHTP01031518.1 | nitrate regulatory gene2 protein-like | -2.51 | 0.00E+00 |
| GHTP01031519.1 | bZIP domain class transcription factor (DUF630 and DUF632) | -2.03 | 1.67E-05 |
| GHTP01032288.1 | small glutamine-rich tetratricopeptide repeat-containing protein | -2.29 | 0.00E+00 |
| GHTP01032810.1 | RING/FYVE/PHD zinc finger superfamily protein | 2.26 | 0 |
| GHTP01033051.1 | protein NRT1/ PTR FAMILY 2.11-like | -5.69 | 2.69E-09 |
| GHTP01034523.1 | arginine/serine-rich coiled-coil protein 2 isoform X2 | -2.85 | 0.00E+00 |
| GHTP01035440.1 | nitrate regulatory gene2 protein | -2.24 | 0.00E+00 |
| GHTP01035776.1 | arginine/serine-rich coiled-coil protein 2 isoform X2 | -2.93 | 0.00E+00 |
| GHTP01037654.1 | ABSCISIC ACID-INSENSITIVE 5-like protein 5 | -3.48 | 0.00E+00 |
| GHTP01037730.1 | probable polyamine oxidase 2 isoform X2 | 2.59 | 0.000242 |
| GHTP01037731.1 | probable polyamine oxidase 2 isoform X2 | 61.64 | 2.12E-13 |
| GHTP01037732.1 | putative polyamine oxidase 2 | 3.44 | 1.38E-08 |
| GHTP01037737.1 | ferredoxin-dependent glutamate synthase. chloroplastic | 4.29 | 0.00E+00 |
| GHTP01039007.1 | AP2-like ethylene-responsive transcription factor ANT | -2.62 | 4.95E-08 |
| GHTP01039235.1 | E3 ubiquitin ligase PARAQUAT TOLERANCE 3-like isoform X1 | -2 | 7.49E-08 |
| GHTP01040150.1 | Tetratricopeptide repeat-like superfamily protein isoform 1 | 2.1 | 0 |
| GHTP01044624.1 | nitrate regulatory gene2 protein-like | -2.59 | 1.57E-05 |
| GHTP01044837.1 | RmlC-like cupins superfamily protein | 2.12 | 3.01E-13 |
| GHTP01045183.1 | nitrate regulatory gene2 protein | -2.06 | 7.13E-04 |
| GHTP01046972.1 | pathogenesis-related protein STH-21-like | 2.69 | 0.00E+00 |
| GHTP01047139.1 | nitrate regulatory gene2 protein-like | -2.12 | 9.01E-09 |
| GHTP01047685.1 | serine/arginine-rich SC35-like splicing factor SCL33 | 2.41 | 8.44E-12 |
| GHTP01048254.1 | fasciclin-like arabinogalactan protein 4 | 5.96 | 2.12E-08 |
| GHTP01048472.1 | proline transporter 2-like isoform X2 | 2.35 | 3.70E-05 |
| GHTP01048773.1 | protein EARLY FLOWERING 3 | 2.16 | 2.19E-09 |
| GHTP01049483.1 | putative S-adenosyl-L-methionine-dependent methyltransferase | 4.08 | 2.01E-10 |
| GHTP01050141.1 | ethylene-responsive transcription factor ABR1-like | 62.09 | 0.00E+00 |
| GHTP01052236.1 | ethylene-responsive transcription factor CRF2-like | 28 | 6.59E-07 |
| GHTP01055544.1 | ethylene-responsive transcription factor ERF073-like | 2.03 | 1.21E-06 |
| GHTP01055617.1 | ethylene-responsive transcription factor ABR1-like | 7.38 | 0.00E+00 |
| GHTP01055971.1 | ninja-family protein AFP2-like | -4.24 | 6.72E-04 |
| GHTP01056931.1 | ethylene-responsive transcription factor ERF027-like | 124.44 | 0.00E+00 |
| GHTP01059070.1 | ABA responsive element binding factor | 4.2 | 2.57E-06 |
| GHTP01062309.1 | ethylene-responsive transcription factor ERF003-like | 7.07 | 8.07E-04 |
| GHTP01063025.1 | precursor of CEP14 | 6.46 | 0.00E+00 |
| GHTP01064836.1 | protein EIN4 | 3.15 | 3.81E-09 |
| GHTP01074812.1 | ethylene-responsive transcription factor ERF054-like | 20.88 | 1.98E-12 |
| GHTP01084431.1 | ethylene-responsive transcription factor ERF027-like | 24 | 4.75E-06 |

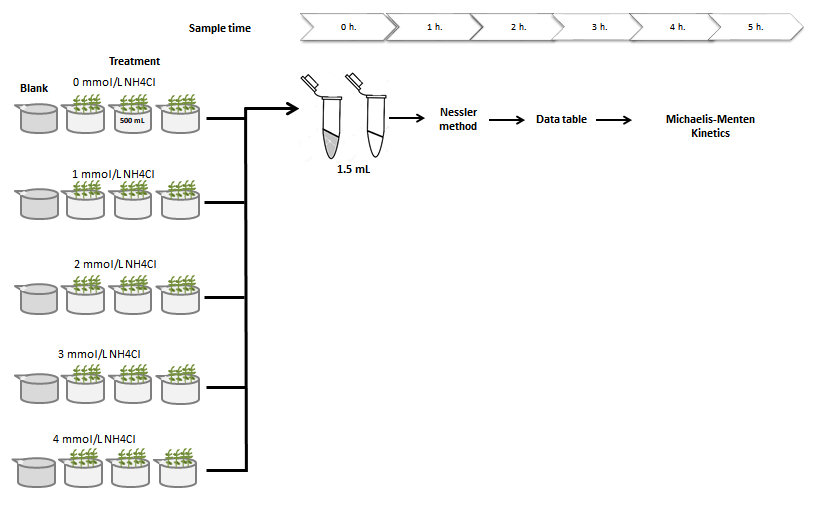

**Fig. S1**. Flowchart of the experimental setup used to collect solution samples for quantitative measurements of changes in ammonium kinetics.

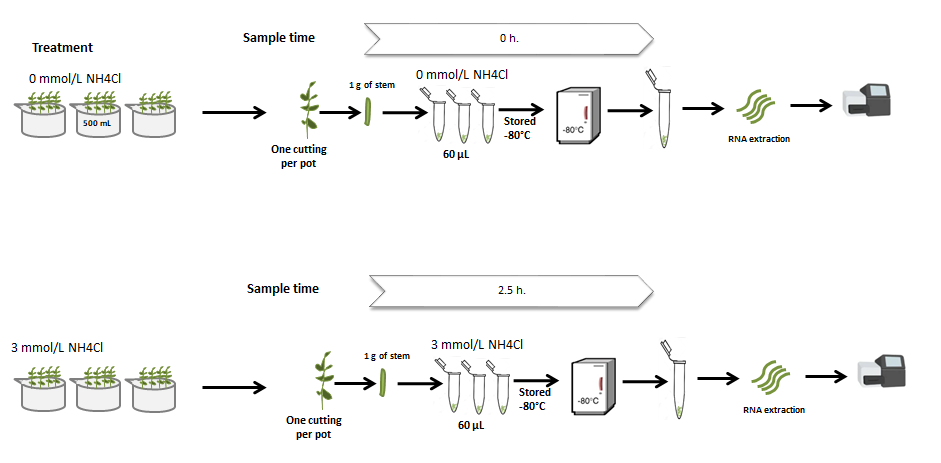

**Fig. S2.** Flowchart of the experimental setup used to collect *S. neei* samples for sequencing.

BlastX against NR database

Match: 45,327

BlastN against *Salicornia*

Match: 32,609

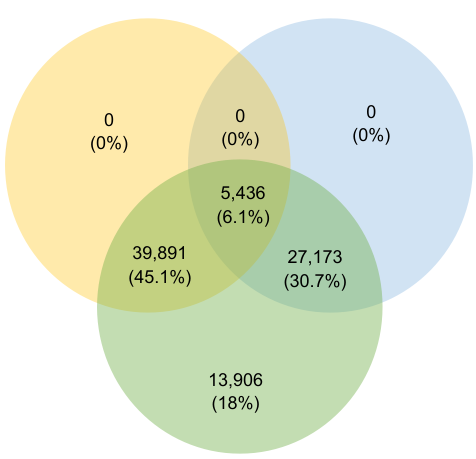

*Salicornia neei*

86,020 contigs

**Fig. S3**. Venn diagram of contigs annotated of *S. neei* with significant blast hits against NR database (orange), NT − Salicornia sequences (blue) and new sequences from *S. neei* that did not match with other databases (green).

In this repository you will find the data and codes to partially reproduce the research carried out on the manuscript.

<https://github.com/GenomicsLaboratory/ReproducibleResearch/tree/main/Diaz_etal_2021>
